# Daam1 negatively regulates USP10 activity

**DOI:** 10.1101/2022.02.14.480271

**Authors:** Andrew T. Phillips, Edward F. Boumil, Arunkumar Venkatesan, Nileyma Castro, Audrey M. Bernstein

**Author notes:** These authors contributed equally (Andrew T. Phillips, Edward F. Boumil). Address correspondence to: Dr. Audrey Bernstein, SUNY Upstate Medical University, Department of Ophthalmology, 505 Irving Ave, Syracuse, NY 13210.

## Abstract

The differentiation of fibroblasts into pathological myofibroblasts during wound healing is in part characterized by increased cell surface expression of αv-integrins. Our previous studies found that the deubiquitinase (DUB) USP10 removes ubiquitin from αv-integrins, leading to cell surface integrin accumulation, subsequent TGFβ1 activation, and myofibroblast differentiation. In this study, a yeast-two hybrid screen elucidated a novel binding partner for USP10, the formin, Daam1. The USP10/Daam1 interaction was also supported by proximity ligation assay (PLA) activity. Treatment with TGFβ1 significantly increased USP10 and Daam1 protein expression, PLA signal, and co-localization to actin stress fibers. Furthermore, Daam1 siRNA knockdown significantly reduced a) co-precipitation of USP10 and Daam1 on purified actin stress fibers, and b) β1- and β5-integrin ubiquitination resulting in increased αv-, β1-, and β5-integrin total protein levels, αv integrin recycling to the cell surface, and extracellular fibronectin (FN) organization. Together, our data suggest that Daam1 negatively regulates USP10’s DUB activity and subsequently maintains integrin protein homeostasis.

## Introduction

Corneal tissue has been used extensively as a model system to study scarring as it is transparent and thus scars in the cornea lead to visual disability and blindness (Stepp et al., 2014; Whitcher et al., 2001). Given the readily available access to non-transplantable donor tissue, primary human corneal cells (HCFs) are used as a model system to discover new pathways that lead to scarring outcomes and these results may be generalized to scarring in other tissues (Bukowiecki et al., 2017; Wilson et al., 2017). Corneal myofibroblasts arise from resident corneal stromal cells termed keratocytes, circulating, bone marrow-derived fibrocytes which readily infiltrate the cornea, as well as through the epithelial-mesenchymal transition of cells within the site of injury (Lassance et al., 2018; Shu and Lovicu, 2017). These cells are important to the wound healing process and normally will have apoptosed in a regeneratively healed wound. However, scar tissue is characterized by the persistence of pathological myofibroblasts (Desmouliere et al., 1995; Wilson, 2012). Myofibroblasts have pronounced actin stress fiber cytoskeleton, characterized by organized α-smooth muscle actin (α-SMA). These prominent stress fibers facilitate myofibroblast-mediated contraction of extracellular matrix (ECM), important in the process of ECM remodeling during wound healing. However, persistent myofibroblasts overly contact tissue, secrete fibrotic extracellular matrix proteins, and promote an autocrine loop of TGFβ activity (Lorenzo-Martin et al., 2019; Massoudi et al., 2016; Pakshir and Hinz, 2018; Sandbo and Dulin, 2011), creating scar tissue.

Integrins also play important roles in wound healing, physically coupling the ECM to the intracellular actin network through focal adhesion complex formation, as well as initiating signaling cascades that induce actin stress fiber contraction and rearrangement in cytoskeletal architecture (Schiller and Fassler, 2013; Thannickal et al., 2003). In regard to fibrosis, the αv-integrins (αvβ1, αvβ3, αvβ5, αvβ6, and αvβ8) are essential to myofibroblast differentiation (Asano et al., 2006; Henderson et al., 2013; Lygoe et al., 2004), with particular αv-heterodimers having more critical roles than others in differing forms of fibrosis (Chang et al., 2017; Horan et al., 2008; Reed et al., 2015; Sarrazy et al., 2014). Our work demonstrated that αvβ1 and especially αvβ5 are most strongly-associated with corneal scarring (Gillespie et al., 2017; Wang et al., 2012). While direct inhibition or knockdown of αv-integrins sequesters myofibroblast differentiation and fibrosis (Chang et al., 2017; Sarrazy et al., 2014), it may also compromise wound healing by interfering with re-epithelialization (Blanco-Mezquita et al., 2011; Duperret et al., 2016).

Previous work by our group identified an alternate strategy for targeting αv-integrins and preventing myofibroblast-induced scar tissue formation. We found that the deubiquitinase (DUB), USP10, is upregulated during myofibroblast differentiation. USP10 deubiquitinates the β1 and β5 subunits of αv-integrin heterodimers, protecting them from degradation and leading to an increase in cell surface expression of αv-dimers. In addition, overexpression of USP10 in HCFs was sufficient to increase the activation of local TGFβ1, induce α-SMA-positive stress fiber formation, and lead to an increase in cellular FN accumulation (Gillespie et al., 2017). In contrast, siRNA knockdown of USP10 significantly reduced the formation of scar tissue in an organ culture wound healing model (Castro et al., 2019; Gillespie et al., 2017), and in vivo in rabbits (Boumil et al., 2020) suggesting that USP10 represents a novel target for intervention in scarring outcomes and perhaps more broadly fibrotic diseases.

The present study further elucidates the regulation of USP10-induced αv-integrin accumulation in myofibroblasts. A yeast 2-hybrid screen, using USP10 as bait, identified novel protein-protein interactions with USP10. Daam1 (Diaphanous-associated activator of morphogenesis 1) was the strongest confidence novel interacting protein with USP10. Daam1 is a formin protein with a diverse set of functions within cells. Formins, including Daam1, are responsible for actin cytoarchitectural arrangement, facilitating such functions as actin filament polymerization (Yang et al., 2007) and severing (Harris et al., 2004), as well as cross-linking actin into bundles (Esue et al., 2008; Jaiswal et al., 2013). Our data suggest that Daam1 negatively regulates USP10, by sequestering the complex to actin stress fibers. Overall, we found that the Daam1/USP10 axis is a novel post-translational regulator of integrin protein levels. These results provide insights into an additional layer of control over the integrin life-cycle.

## Results

### Yeast 2-hybrid screen reveals novel Daam1-USP10 protein-protein interaction

In order to further characterize the mechanisms by which USP10 promotes integrin recycling and aids in the differentiation of myofibroblasts, a yeast 2-hybrid screen (Hybrigenics) was carried out using USP10 as bait, to identify novel targets for study. The screen produced both established and novel hits (Fig 1A). The established hits included both G3BP proteins; G3BP stress granule assembly factor 1 (G3BP1; NCBI Ref # NM_005754.2) and G3BP stress granule assembly factor 2 (G3BP2; NCBI Ref # NM_203504 and NM_012297). These were considered strong positive controls for this screen. The strongest confidence novel interaction identified by our yeast 2-hybrid screen for USP10 binding partners was with the formin protein Disheveled-associated activator of morphogenesis 1 (Daam1; NCBI Ref # NM_001270520.1), the primary subject of this study. This interaction was denoted as “Very Strong” confidence, as determined by a proprietary calculation developed by Hybrigenics to screen out putative false positive interactions. Of note was the USP10 interaction signal from the protein Fascin-1 (NCBI Ref # NM_003088), another actin-associated protein that directly interacts with Daam1 on actin bundles in lamellipodia (Jaiswal et al., 2013). These are thought to be structural interactions as using USP10 as bait did not reveal known enzymatic interactions in which USP10 acts as a DUB such as p53 or integrins (Gillespie et al., 2017; Yuan et al., 2010).

**Figure 1.**
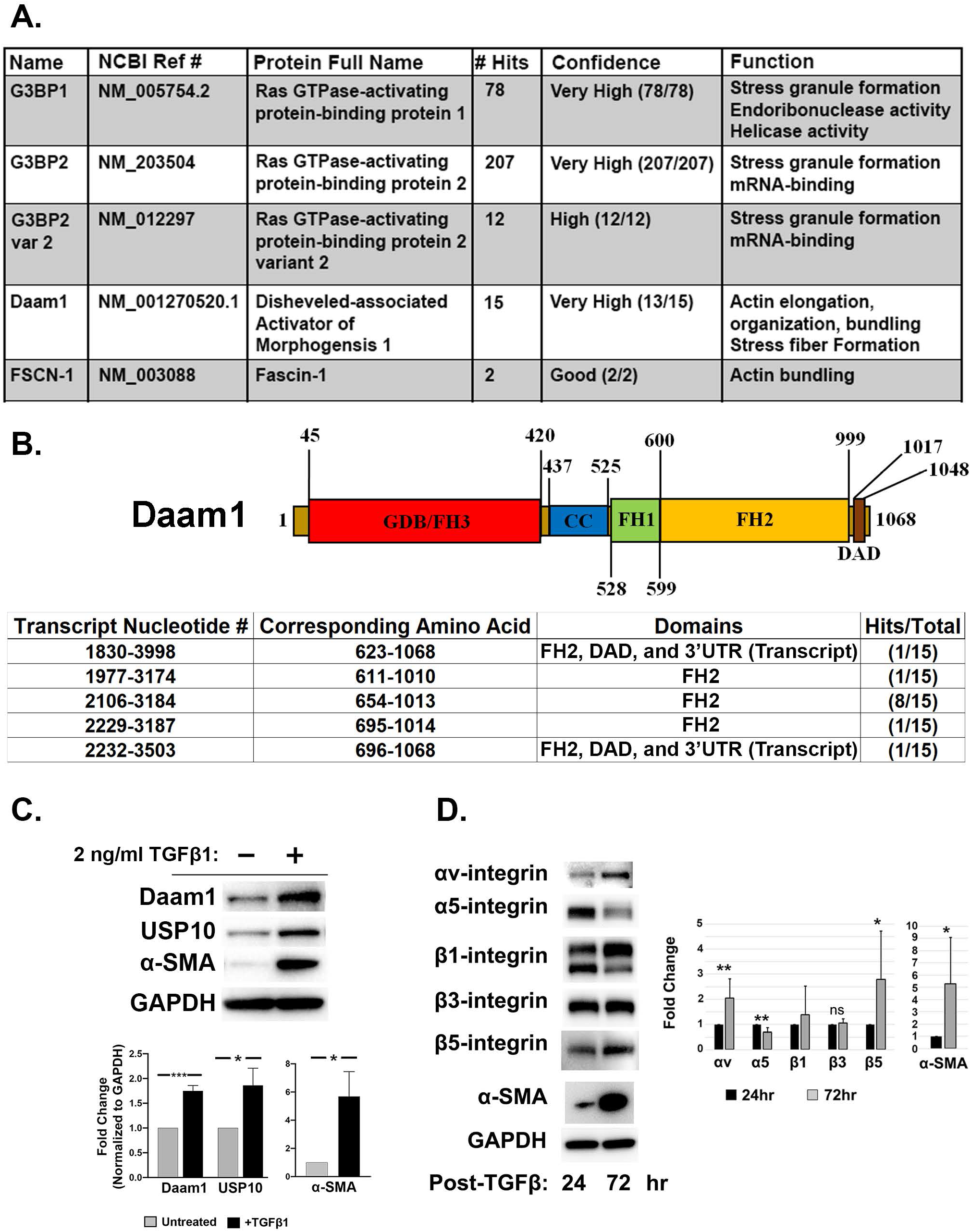
Yeast 2-Hybrid screen reveals Daam1 as a novel USP10-binding partner. A) Yeast 2-hybrid data (Hybrigenics), human placental mRNA library, and human USP10 as bait. B) Daam1 subdomain schematic and yeast 2-hybrid fragment data. C) Immunoblot with quantification of untreated and TGFβ1 treated HCFs (72 hrs). Quantification is expressed as the mean fold change after normalization to GAPDH. D) Comparison of HCF lysates treated for 24 and 72hrs with TGFβ1. N=3. Statistical significance was calculated using an unpaired t-test +/- SEM.

Daam1 is composed of 4 primary functional domains: GTPase-Binding Domain/Formin Homology Domain 3 (GBD/FH3), Formin Homology Domain 2 (FH2), Formin Homology Domain 1 (FH1), and the Diaphanous Autoregulatory Domain (DAD), as well as a coiled-coil (CC) structural domain between the GBD/FH3 domain and the FH2 domain. FH2 domain was obtained as the site of interaction in all readable hits (Fig 1B). The yeast 2-hybrid screen provides information about the fragment of the mRNA which generates the interaction. Of the 15 hits for Daam1, 12 gave readable sequence data. Eight hits were specific to a fragment of the parent Daam1 mRNA containing only the formin homology domain 2 (FH2), while 2 fragments were specific to the FH2 domain and the C-terminal diaphanous auto-regulatory domain. The remaining 3 fragments were not capable of providing sequence related information.

Next, we determined if Daam1 is expressed in HCFs by western blot. Daam1 was detected in lysates from cultured HCFs at a characteristic 123 kDa band, confirming its expression by these cells (Fig 1C). In addition, incubation of HCFs in 2ng/mL of TGFβ1 for 3 days significantly increased Daam1 expression by 1.8-fold (p<0.001), a similar result to what was observed in myofibroblasts in lung samples of idiopathic pulmonary arterial hypertension patients (Yanai et al., 2017), as well as total tissue gene expression analysis of kidney samples following injury (Saito et al., 2015). TGFβ1-induced differentiation of HCFs into myofibroblasts was confirmed by a well-characterized increased in expression of α-SMA (4.7-fold p<0.05). Our previously-published finding that USP10 expression increases in myofibroblasts was also found to be reproducible (1.9-fold increase, p<0.05; (Gillespie et al., 2017)).

Finally, because our model system includes treatment of HCFs with TGFβ1 to enhance protein expression and elucidate protein interactions, we investigated the effect of TGFβ1 treatment of integrin expression in HCFs over time (between 24 and 72hr; Fig 1D). Expression of αv, β5, α-SMA were all found to significantly increase over this time course (2.1-fold, p<0.01; 2.8-fold, p<0.05; and 5.3-fold, p<0.05, respectively), whereas β1 trended towards increase but was not significant. β3-integrin expression did not change over this same time, as has been previously observed (Gillespie et al., 2017). Interestingly, α5-integrin levels were reduced at 72hrs by 32% (p<0.01).

### TGFβ1 treatment promotes Daam1-USP10 Proximity

To support the results of the yeast 2-hybrid screen, proximity ligation assay (PLA) was carried out in cultured HCFs either untreated or treated with 2ng/ml TGFβ1 for 3 days (Fig 2). After optimization of assay conditions, positive signal between USP10 and Daam1 was achieved in untreated cells, which appeared as bright, randomly dispersed puncta present through the cytoplasmic space (Fig 2A). Puncta are defined as a positive indicator of interaction of target proteins with a spatial resolution of <30-40 nm (Zatloukal et al., 2014). Signal was quantified by puncta density to analyze the change in interaction frequency. The USP10/Daam1 PLA pair increased in signal density between untreated and TGFβ1 treated cells by 1.2-fold (*p*<0.05) (Fig 2B). PLA signal was significantly reduced by omission of either primary antibody, suggesting the signal was not due to nonspecific interaction by oligo-conjugated secondary antibodies. The USP10/G3BP2 PLA pair in cells treated with TGFβ1 (which served as a positive control for the assay) produced the highest signal, with a 1.4-fold (*p*<0.001) higher puncta density than the USP10/Daam1 PLA signal without TGFβ treatment and 1.2-fold (*p*<0.05) higher density than the USP10/Daam1 PLA signal in cells treated with TGFβ. Overall, these data demonstrate that USP10 is in proximity to both Daam1 and G3BP2 and that TGFβ increases the USP10/Daam1 interaction.

**Figure 2.**
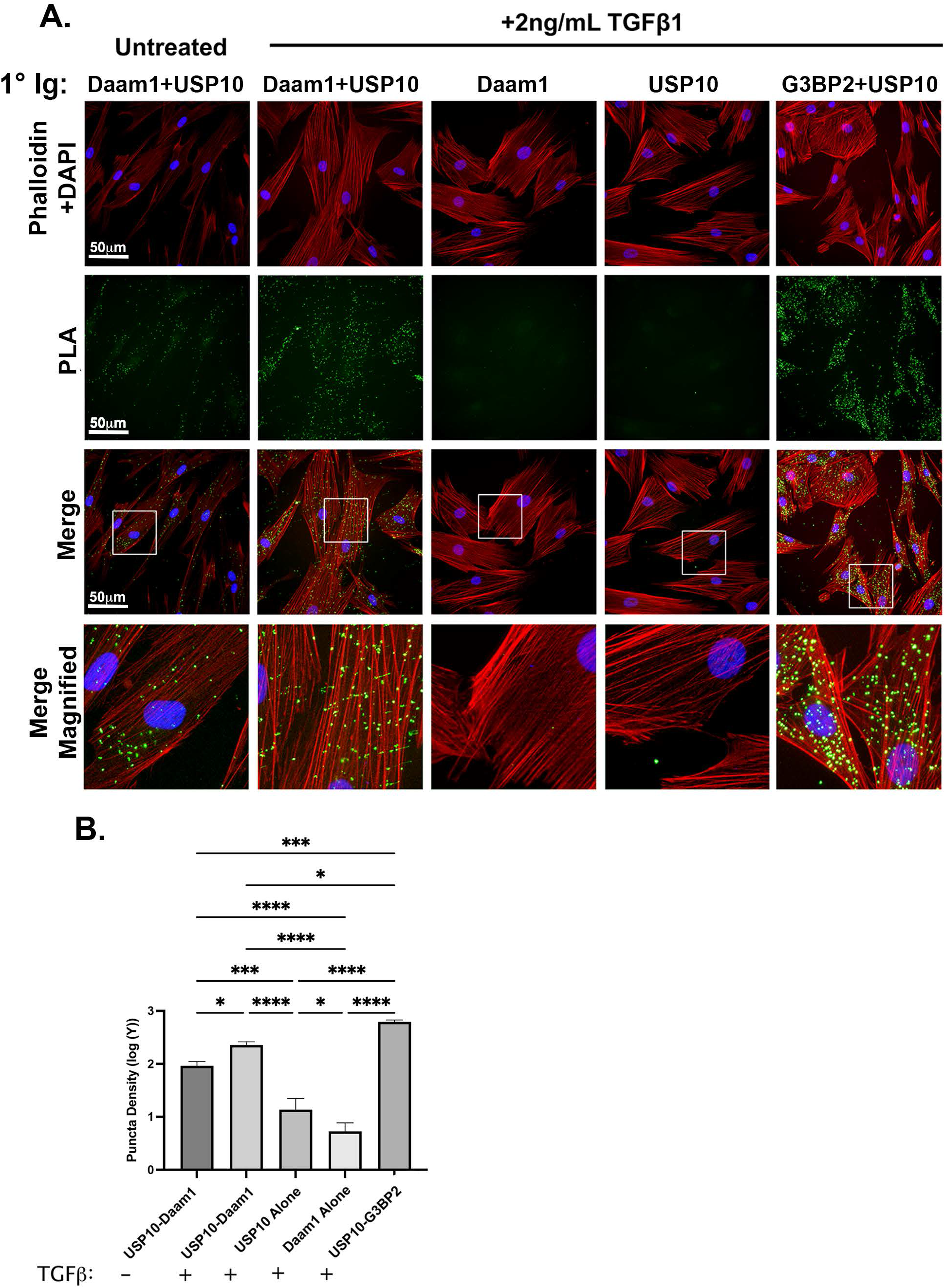
Proximity Ligation Assay supports USP10-Daam1 interaction in HCFs. A) Representative images of untreated and TGFβ1 treated HCFs (PLA, green), counterstained for actin (phalloidin, red) and nucleus (DAPI, blue). B) Quantification of puncta density in each condition. Stroked portions of the merged channels are magnified (Merge Magnified). Bar=50μm. A total of 15 cells were analyzed for each condition from 3 independent experiments. Statistical significance was calculated using one-way ANOVA after log transformation of the data +/- SEM.

### TGFβ1 drives USP10 and Daam1 localization to actin stress fibers

To determine the localization of Daam1 and USP10 in HCFs, cells were cultured without or with TGFβ1. Untreated HCFs immunostained with Daam1 antibody resulted in a diffuse and randomly-organized punctate fashion, scattered throughout the cell (Fig 3A). Following TGFβ1 treatment, as expected α-SMA antibody labeled cells demonstrated a characteristic filamentous stress fiber phenotype. Daam1 likewise exhibited a “ribbed” or filamentous pattern that corresponded with the presence of α-SMA stress fibers in cells. Omission of Daam1 primary antibody abolished 488-fluorescence, suggesting staining is not due to non-specific secondary antibody labeling.

**Figure 3.**
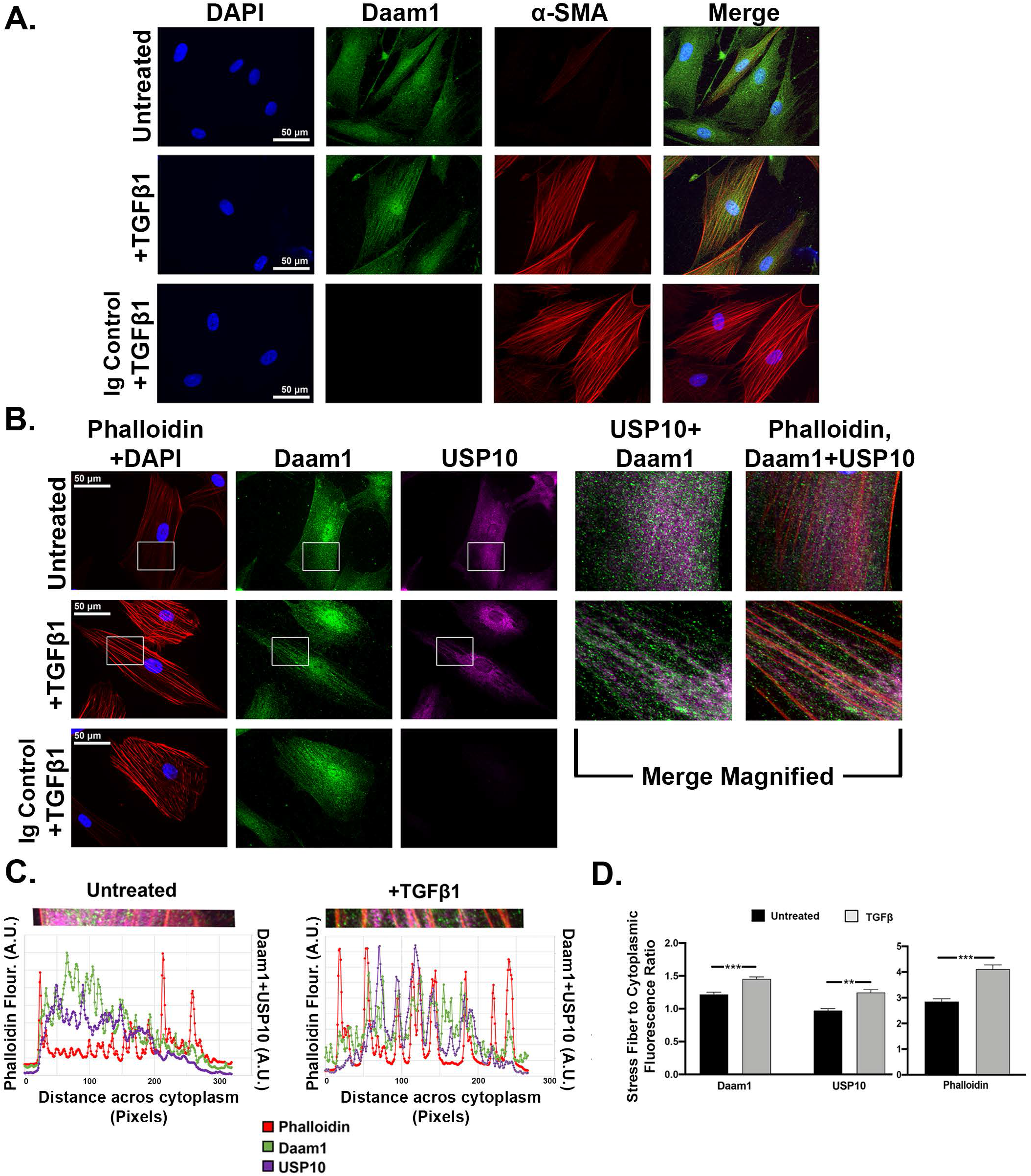
TGFβ1 Drives Daam1 and USP10 Localization Toward Actin Stress Fibers. A) Analysis of untreated and TGFβ1-treated HCFs, with DAPI (blue), α-SMA (red), and Daam1 (green) immunostaining. TGFβ1 treatment drives Daam1 immunofluorescence from a diffuse, punctate distribution to a stress fiber pattern. IgG control abolished immunofluorescence signal. B) Immunostaining for USP10 (violet), phalloidin (red), Daam1 (green). Like Daam1, TGFβ1 treatment drives USP10 immunofluorescence from a diffuse pattern to a stress fiber pattern. C) Fluorescence intensity profiling across representative cells (from panel B) in scatterplot format. D) Quantification of multiple fluorescence intensity profiles, comparing untreated (black) and TGFβ1-treated cells (grey), expressed as a ratio of fluorescence intensity at stress fibers to the intensity between stress fibers (cytoplasmic). An increase in stress fiber association with TGFβ1 was observed for all three proteins. Statistical significance was calculated using an unpaired t-test +/- SEM. Bar= 50μm. N=3

Treated and untreated HCFs were then immuno-labelled with USP10 antibody (with phalloidin used as a counter stain because of IgG host restrictions; Fig 3B). Like Daam1, USP10 was observed in a diffuse manner throughout the cytoplasm of untreated HCFs, with a slight perinuclear bias in intensity. Following TGFβ1 treatment, USP10 also underwent a shift in localization, with an increased coincidence with actin stress fibers.

We quantified this transition of USP10 and Daam1 fluorescence by measuring fluorescence intensity profiles across the perinuclear cytoplasm (Fig 3C), defining a threshold fluorescence level which segregated the cytoplasmic space from actin stress fibers as labeled by phalloidin, and then binning the corresponding fluorescence intensities in the Daam1and USP10 channels into either “Cytoplasmic” or “Stress Fiber” group and averaging them. The Stress Fiber bin was divided by the Cytoplasmic bin to generate a Stress Fiber-to-Cytoplasmic intensity ratio for each cell, and averaged across all cells for statistical analysis (Fig 3D). We predicted that a random distribution of Daam1 and USP10 would result in Stress Fiber-to-Cytoplasmic ratios with values closer to 1 (meaning equal average intensity associated with stress fibers and cytoplasm), and that TGFβ1 would increase these values based on our observations. Daam1 stress fiber-to-cytoplasmic ratio was 1.22 in untreated cells, TGFβ1 treatment increased this ratio to 1.45 (p<0.001). Thus, Daam1 transitioned from a distribution that was, on average, 1.22-fold higher at stress fibers than in cytoplasmic areas, to a distribution where Daam1 intensity was 1.45-fold higher at stress fibers than in the cytoplasm. USP10 exhibited a similar shift, changing from 0.97 in untreated cells to 1.25 in TGFβ1 treated cells (p<0.001). Phalloidin, which by definition should be much greater than 1 as it was used to label stress fibers, likewise experienced a shift of 2.85 to 4.11 (p<0.001).

### Daam1 siRNA reduces Daam1 and USP10 binding to stress fibers

In order to confirm the observation from the immunocytochemical analyses that Daam1 and USP10 associate with actin stress fibers, we utilized a protocol to isolate intact stress fibers from fibroblasts (Fig 4). Our hypothesis was two-fold: 1) if Daam1 and USP10 stress fiber localization is due to physical cytoskeleton-associated complex formation, both proteins should be coprecipitated with actin stress fibers, and 2) if Daam1 (a canonical actin binding protein) recruits USP10 to stress fibers, Daam1 knockdown will result in a concomitant reduction in USP10-stress fiber coprecipitation.

**Figure 4.**
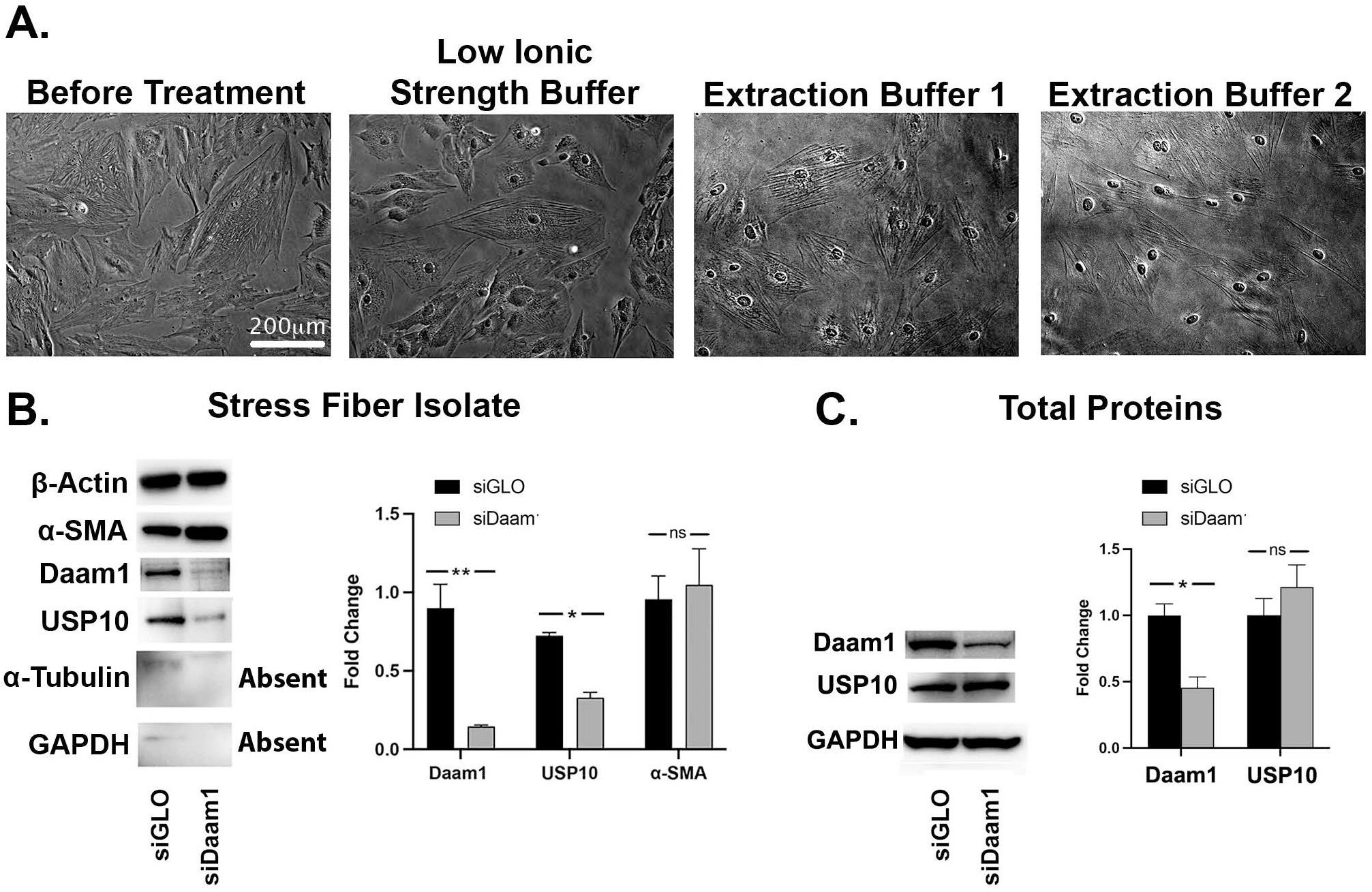
Daam1 and USP10 precipitate with stress fibers. A) Representative phase-contrast images of TGFβ1 treated HCFs subjected to a series of buffer washes with increasing concentration of detergent. Bar=200μm B) Immunoblot analysis of stress fiber isolates generated from TGFβ1 treated cells transfected with nontargeting siGLO or siDaam1. Daam1 knockdown also reduced USP10 association with stress fibers. C) Immunoblot analysis of total protein from HCFs transfected with siGLO or siDaam1. siDaam1 did not significantly affect total USP10 levels. Statistical significance was calculated using an unpaired t-test +/- SEM. N=3

HCFs were transfected with either non-RISC associating siRNA (siGLO) or a pool of siRNAs against human Daam1 (siDaam1), and cultured with TGFβ1 for 3 days (to promote stress fiber formation). Images were taken of the cultures as they underwent washes in buffers with increasingly strong detergents to monitor extraction progress (Fig 4A). Cells became increasingly difficult to observe by phase contrast microscopy as the extraction process progressed, suggesting a decrease in cell density as non-stress fiber associated components were stripped away. Stress fibers remained visible and intact throughout the process, indicating proper preservation of this cytoskeletal element. Following collection of the stress fibers and enrichment by ultracentrifugation, precipitates were processed and analyzed by western blot (Fig 4B). Quantification is expressed as fold-changes after normalization to β-actin. The precipitates from both siGLO and siDaam1 transfections were rich in both β-actin and α-SMA, which developed with ≤1 second exposure times, suggesting proper enrichment of the actin cytoskeleton. siDaam1 did not induce any significant changes in α-SMA levels as compared to siGLO (*p*>0.05). Conversely, the normally heavily abundant proteins GAPDH and α-tubulin was barely (or not at all) detectable, even with abnormally long exposure times (>30 sec, as compared to ~1 sec for total cell lysates in previous experiments). Precipitates from siGLO transfected cultures stained strongly for both Daam1 and USP10, suggesting they are physically associated with stress fibers. Transfection of siDaam1 reduced Daam1 co-precipitation with stress fibers by 87.0% (*p*<0.001). Further, siDaam1 transfection also resulted in a 60.1% reduction in USP10 coprecipitation with stress fibers (*p*<0.001). siDaam1 treatment did not significantly affect total USP10 levels in unfractionated lysates from HCFs (Fig 4C; *p*>0.05). These data suggest that Daam1 may either recruit USP10 to stress fibers or is at least in part necessary for its association.

### Daam1 knockdown promotes integrin protein accumulation with reduced integrin ubiquitination

Because TGFβ is a fibrotic growth factor, and we know from our work and others that USP10 and Daam1 are increased under fibrotic conditions, we reasoned that Daam1 was a positive regulator of USP10’s DUB activity. However, in the following experiments, we found that the opposite result, our data support the idea that Daam1 negatively regulates USP10’s DUB activity.

To assess the biological function of USP10-Daam1 interaction in HCFs, we analyzed integrin subunit protein levels without or with Daam1 knockdown by immunoblot analysis (Fig 5A), with quantification of densitometry normalized to GAPDH loading controls. In this experiment we continued to use TGFβ1 treatment of HCFs to increase total integrin protein levels. siDaam1 transfection resulted in a 69.8% reduction in Daam1 levels vs siGLO (*p*<0.001). Daam1 knockdown increased total αv-, β1-, and β5-integrin levels in HCFs by 1.6-fold, 1.4-fold, and 1.4-fold, respectively, vs. siGLO controls (*p*<0.05 for each). Daam1 knockdown also elevated α5-integrin levels by 1.3-fold, however this change was not significant (*p*=0.065), and no effect was observed on β3-integrin levels, as had been seen previously on USP10’s effect on β3-integrin (Gillespie et al., 2017). Collectively, these results demonstrate that Daam1 regulates total levels of αv, β1, and β5 integrin protein expression. These data support the idea that Daam1 is negatively regulating USP10’s DUB activity on integrins. We propose that knocking down Daam1 releases USP10 activity, leads to less ubiquitination and degradation of integrins, and therefore an increase in total integrin protein.

**Figure 5.**
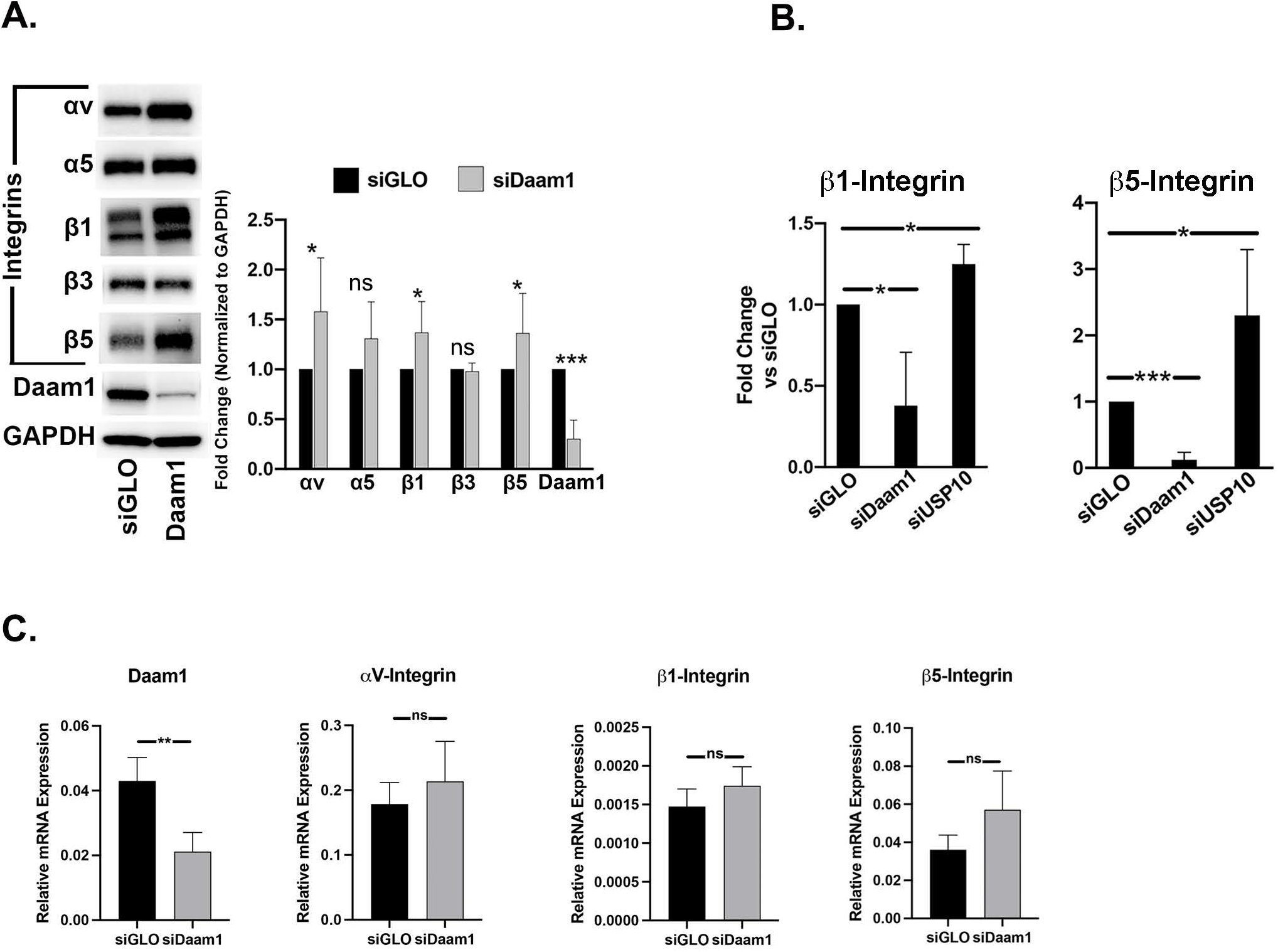
Effect of Daam1 knockdown on integrins. A) HCFs were transfected with siDaam1 or siGLO and treated with TGFβ for 3 days to increase integrin expression. siDaam1 transfection resulted in increased expression of αv-, β1-, β5-integrins. N=6. B) HCFs were transfected with siDaam1 or siGLO. After 3 days cells were lysed and subjected to Ubiquant™ ubiquitin capture ELISA. siDaam1 reduced, whereas siUSP10 increased ubiquitination of β1 and β5. N=4. (C) HCFs were transfected with siGLO or siDaam1 and treated with TGFβ for 3 days prior to processing for qPCR. The relative expression of Daam1, integrin αV, β1, and β5 is shown as compared to GAPDH. Statistical significance was calculated using an unpaired t-test Bar=200μm. N=3.

To test this, we utilized a sensitive ubiquitin capture ELISA (Ubiquant™), with this assay, treatment with TGFβ was not necessary to obtain significant results. HCFs were transfected with either siGLO, siDaam1 siRNA, or as a positive control, human USP10 targeting siRNA (siUSP10) (Gillespie et al., 2017), and cultured for 3 days (Fig 5B). Compared to siGLO, siDaam1 transfection resulted in a 62.1% decrease in β1-integrin ubiquitination (*p*<0.05), conversely, siUSP10 increased β1-integrin ubiquitination by 1.2-fold (*p*<0.05). Similarly, daam1 knockdown decreased β5-integrin ubiquitination by 87.9% (*p*<0.001), whereas USP10 knockdown increased β5-integrin ubiquitination by 1.3-fold (*p*<0.05). The results with USP10 siRNA mirrored our previous published results (Gillespie et al., 2017). These data suggest that less Daam1 expression (siDaam1) leads to increased USP10 activity (removes more ubiquitin) and reduced integrin ubiquitination. Knockdown of USP10 (siUSP10) has the opposite effect. Together our data suggest that Daam1 is a negative regulator of USP10’s DUB activity on integrins. To test the effect of siDaam1 on integrin gene expression, HCFs were transfected with siGLO or siDaam1 and treated with TGFβ for 3 days prior to processing for qPCR. The relative expression of Daam1, integrin αV, β1, and β5 is shown as compared to GAPDH. Whereas Daam1 knocked down by 50.7% (*p*<0.01), the other relationships were not significant (Fig 5C), supporting the idea that the effect of Daam1 on integrins is post-translational.

### Daam1 knockdown results in αv integrin and FN cell surface accumulation

To further link the Daam1/USP10 interaction to integrins, we tested if Daam1 knockdown would increase integrin recycling, and a functional consequence of increase integrin recycling, FN recycling. For the integrin recycling assay, 48 hrs post-transfection cells were blocked and treated with Ab for 30 minutes prior to cell surface stripping for 30 sec. Cells were futher incubated for 90 min prior to incubation with 2° Ab-488 for 30 min (Fig 6A). Using live cell confocal microscopy, cell surface integrin signal was imaged. In parallel with our western blot findings, whereas there was a significant increase in αv integrin recycling (2.1 fold *p*<0.05), α5β1 integrin recycling was not significantly different between conditions (Fig 6B,C).

**Figure 6.**
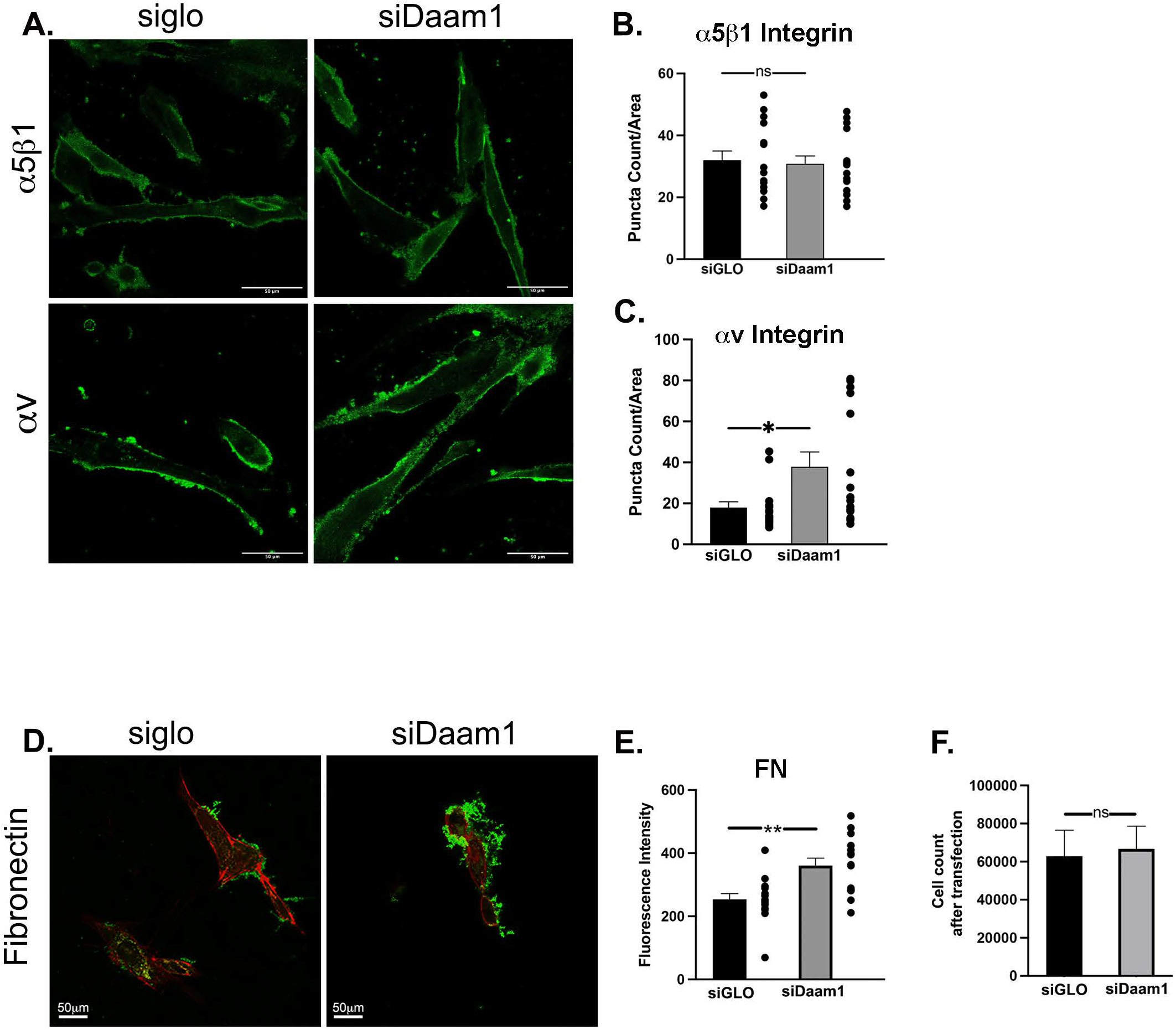
Knockdown of Daam1 increases integrin and fibronectin (FN) recycling. A) Live cell integrin recycling assay. 48 hrs post-transfection cells were blocked and treated with Ab against α5β1 or αv at 10ug/ml for 30 minutes prior to cell surface stripping for 30 sec. Cells were incubated for 90 min prior to incubation with 2° Ab-488 for 30 min. Bar=50μm. Quantification of recycled (B) Integrin α5β1 and (C) Integrin αV, N=3 with a total of 15 images analyzed for each condition. (D) Live cell FN recycling assay. HCFs were transfected as stated above. 24hrs post transfection HCFs were loaded with biotinylated-FN for 3 hours. After trypsinization (to separate cells from extracellular, non-internalized FN), HCFs were replated and imaged 48 hours post transfection. Prior to imaging, cells were incubated with streptavidin-488 to detect only recycled biotinylated-FN. Imaged by live cell confocal. Bar= 50μm. N=3. D) Quantification of extracellular FN. A total of 15 cells were analyzed for each condition from 3 independent experiments. E) Cell numbers 2 days after transfection for each condition are not significantly different. N=3. Image analysis was done using ImageJ’s Analyze Particles function. Statistical significance was calculated using an unpaired t-test +/- SEM.

Our recent work demonstrates that increased USP10 overexpression that leads to the accumulation of cell surface integrins, (Gillespie et al., 2017) also results in the integrin-mediated accumulation of extracellular FN. (Phillips et al., 2021) Furthermore, that in cell culture, approximately 1/3 of the deposited extracellular FN was derived from recycled FN (Phillips et al., 2021). To determine if Daam1 knockdown affects FN recycling, HCFs were transfected with siGLO or siDaam1. After 24 hours cells were treated with biotinylated-FN for 3 hours, prior to trypsinization to remove uninternalized biotinylated-FN and reseeding. After 48 hours, external FN was detected with Streptavidin-488 (green), and associated cells were detected with SiR-actin (red). In line with an increase in total and recycled integrin, FN accumulation increased significantly in the siDaam1 condition, 1.4 fold p< 0.01 (Fig 6D,E). Fig 6F demonstrates that equal numbers of cells survived the siGLO and siDaam1 transfections.

## Discussion

Integrins are dynamic molecules, and in general cells must have the ability to rapidly fine tune the localization or strength of focal contacts to carry out a diverse array of functions (Huttenlocher and Horwitz, 2011; Schmidt and Friedl, 2010). In order to achieve this, integrins are continuously in a state of flux, internalized into endosomes and either degraded by the endolysosomal system or shuttled back to the cell membrane in a regulated manner (Bridgewater et al., 2012). Our previous work identified USP10 as a critical regulator in the dynamic control of integrins, shifting the balance from intracellular degradation to cell surface accumulation after wounding (Gillespie et al., 2017). The present study further expands on that work by providing an additional point of control on integrin turnover through the novel interaction of USP10 with Daam1.

The formin, Daam1, has been well characterized as an effector of the actin cytoskeleton, influencing such processes as ciliogenesis (Corkins et al., 2019), filopodia extension (Jaiswal et al., 2013), and cell polarity (Ju et al., 2010; Nishimura et al., 2016), to name a few. Daam1 has been found to be upregulated in idiopathic pulmonary fibrosis (IPF) (Rydell-Tormanen et al., 2016), likely due to increased WNT signaling during the wound healing process (Konigshoff et al., 2008; Newman et al., 2016), linking Daam1 to fibrotic conditions. In addition, 4 SNP’s have been identified in GWAS of Caucasian patients with IPF near the Daam1 gene, providing further evidence for a connection to fibrosis (Manichaikul et al., 2017). Furthermore, Daam1 may influence wound healing by promoting haplotaxis (Zhu et al., 2012) whereby epithelial cells and fibroblasts migrate to the site of injury to repopulate and repair a wound (Basan et al., 2013; Blanco-Mezquita et al., 2013).

Lead by a yeast-two hybrid study using USP10 as bait, here we demonstrate a novel role for Daam1 as a binding partner of USP10 (Fig 1A), likely through an interaction with Daam1’s FH2 domain (Fig 1B). Daam1 has numerous binding partners critical to its function, including the integrin-associated kinase Src (Aspenstrom et al., 2006), the Wnt-pathway associated protein Disheveled (Dvl) to the DAD of Daam1, and Rho-GTPase association with the GBD/FH3 subdomain (Liu et al., 2008). However, FH2 domain is canonically thought to directly associate with actin, and to facilitate actin polymerization (Higgs, 2005; Higgs and Peterson, 2005). Thus, our data suggests that Daam1 has a non-canonical role as a part of an USP10-Daam1 FH2 axis, linking Daam1 to integrin turnover and the ubiquitin system. This interaction was supported by proximity ligation assay data (Fig 2). We found that treatment with TGFβ1 enhanced the proximity between USP10 and Daam1 and promoted co-localization with stress fibers (Fig 2 and 3). Daam1 and USP10 localization in the absence of TGFβ1 was observed to be diffuse throughout the cell. USP10’s localization was also perinuclear, an observation we confirm through quantification of the ratio fluorescence associated with stress fibers over the unassociated (cytoplasmic) with stress fibers. One interpretation of this result is that this interaction reflects a spatial aspect of USP10’s function in integrin recycling, perhaps implicating localization to the perinuclear recycling compartment (PNRC). Another possibility is that USP10 associates with early (sorting) endosomes, responsible for facilitating the recycling of a number of membrane proteins, such as CFTR and integrins (Bomberger et al., 2009; Jonker et al., 2018)

The stress fiber localization of USP10 and Daam1 proteins was supported and expanded on by co-precipitation of these proteins during biochemical isolation of actin stress fibers from cells (Fig 4). siDaam1 knockdown reduced Daam1 and USP10 precipitation with stress fibers (Fig 4B), despite having no effect on total USP10 levels (Fig 4C) suggesting that Daam1 may recruit USP10 to stress fibers. siDaam1 treatment did not have a significant effect on α-SMA composition of stress fibers, suggesting changes in USP10 were not due to gross defects in stress fiber assembly. It also may be possible that other actin-associated proteins, such as Fascin-1 (Fig 1A), are in this complex with Daam1, tethering USP10 to actin stress fibers.

From the data in Figs 1–4, we postulated that Daam1 was necessary to “activate” USP10 because TGFβ1 treatment, a typical wounding model, increased association and our previous data showed that wounding and TGFβ1 treatment increased USP10 expression. We hypothesized that Daam1 was structurally necessary for USP10’s DUB activity. However, data in Fig 5 countered that model. Indeed, we found that in fact Daam1 is a “brake” for USP10 negatively regulating its activity. USP10 overexpression post-translationally increases integrin protein levels with no effect on gene expression (Gillespie et al., 2017). USP10 removes ubiquitin from the integrin β1 and β5 subunit and thus USP10 overexpression decreases integrin ubiquitination and degradation and integrins accumulate in the cell (Gillespie et al., 2017). Here, we reasoned that if Daam1 is a brake for USP10, that reducing Daam1 would increase USP10 activity, removing ubiquitin leading to increased protein levels (Fig 5A) and decreased ubiquitination of integrins β1 and β5 (Fig 5B). Daam1 knockdown had no effect on β3-integrin, a result also found during the previous investigation into USP10’s effect on integrin levels (Gillespie et al., 2017). The integrin αv subunit was not tested because it is not ubiquitinated (Hsia et al., 2014; Lobert and Stenmark, 2010). In Fig 5C, we demonstrated that by qPCR after Daam1 knockdown, integrin subunits are not significantly changed, although they are trending towards an increase. These data suggest that the effect of siDaam1 in HCFs is primarily post-translational (through USP10’a DUB activity), however it is possible that there is an indirect effect on gene transcription with siDaam1 (less Daam1 > more USP10 activity > more TGFβ activity) (Gillespie et al., 2017) which eventually leads to an increase in integrin gene expression (Munger and Sheppard, 2011). Taken together, these data (Fig 5) support the hypothesis that Daam1/USP10 interaction is inhibitory to USP10’s ability to deubiquitinate β1- and β5-integrin subunits.

Daam1 serving as a negative regulator of USP10 is further supported by an increase in integrin and FN recycling (Fig 6) after Daam1 knockdown. FN undergoes an extracellular stepwise integrin-dependent polymerization to generate fibrils from soluble, monomeric FN (Mao and Schwarzbauer, 2005; Pankov et al., 2019; Schwarzbauer and Sechler, 1999; Wierzbicka-Patynowski et al., 2004). α5β1 and αv integrins recognize the common integrin-binding motif (RGD) in FN (Benito-Jardon et al., 2020; Danen and Sonnenberg, 2003; Huveneers et al., 2008) and coordinate to achieve efficient FN binding (Benito-Jardon et al., 2021; Bharadwaj et al., 2017). Using a live cell recycling assay, we demonstrated an increase in αv integrin α5β1 integrin recycling, and integrin-mediated FN recycling after USP10 overexpression (Phillips et al., 2021). Here we find that after Daam1 knockdown, similar to USP10 overexpression and in line with integrin accumulation (Fig 5), an increase in αv integrin and extracellular FN recycling (Fig 6). We did not find a significant increase in α5 protein levels (Fig 5A) or α5β1 integrin recycling (Fig 6B) suggesting perhaps that the Daam1/USP10 axis more directly effects αv integrins.

In Fig 7, we propose a model that fits our data. First, although TGFβ is frequently used as a wounding model, in general it upregulates protein expression of many proteins. In this context, both USP10 and Daam1 were upregulated with TGFβ treatment. However, we found that Daam1 is a brake for USP10-mediated integrin protein accumulation, allowing for an additional layer of homeostatic control over total integrin levels in a wounding environment, possibly by creating a USP10 “sink” on stress fibers. TGFβ treatment of HCFs appears to have elucidated this part of the USP10 life cycle.

**Figure 7.**
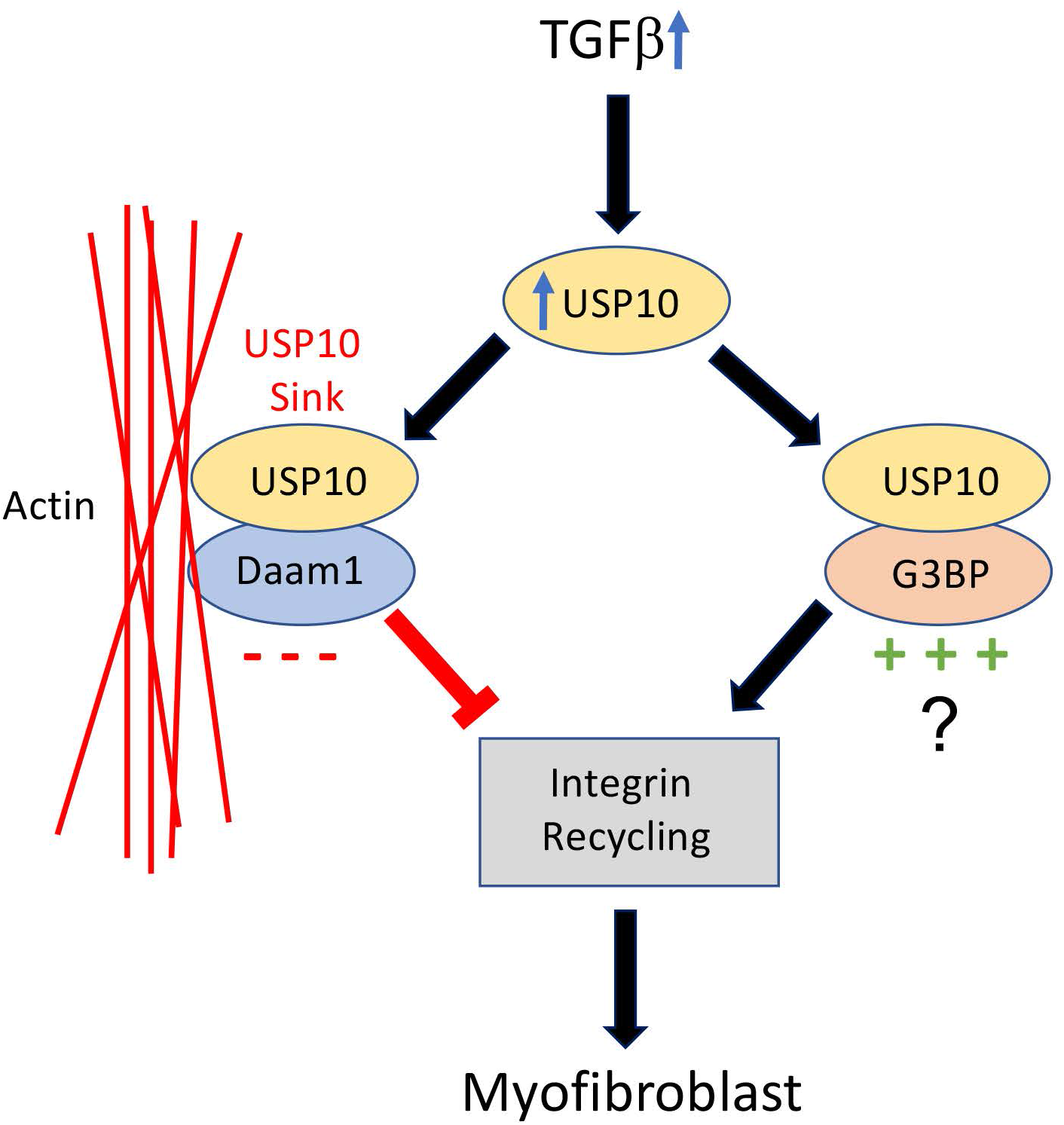
Working model. In response to stress-induced TGFβ1 release, both USP10 and Daam1 expression is increased. Daam1 sequesters USP10 to stress fibers, and USP10’s DUB activity on integrins is inhibited by Daam1. This may act as a level of control over USP10 activity in myofibroblasts. We further hypothesize that the known interaction of USP10 and G3BP1/2 proteins (also shown in Fig 1) positively regulates USP10’s DUB activity on integrins leading to integrin accumulation. The previously known roles of G3BP1/2 in integrin signaling support this concept.

In terms of positive regulators of the USP10/integrin axis, although it is currently unknown, studies indicate that the G3BP proteins could be activators of USP10 activity on integrins. In terms of G3BP/USP10 binding, in addition to our yeast-two hybrid study and PLA (Fig 1A, Fig 2), it was previously demonstrated that USP10 binds to G3PB2 in the cytosol, inducing p53 cytoplasmic localization, ubiquitination, and degradation (Takayama et al., 2018). Connecting G3BP to integrins is the finding that G3BP knockdown inhibits Scr/FAK/ERK signaling, and human lung cancer cell migration and invasion, suggesting a USP10/integrin/G3BP complex and coordination between these proteins (Zhang et al., 2013). Further studies will elucidate these interactions. In summary, our studies indicate that DUBs are a key point of regulation in the integrin life-cycle and may be therapeutic targets for a broad range of integrin-mediated pathologies.

## Materials and Methods

### Cell Culture

Human cadaver corneas from unidentifiable diseased subjects were obtained from the Syracuse Eye Bank, Syracuse, NY and The Eye-Bank for Sight Restoration, New York City, NY. The SUNY Upstate Medical University Institutional Review Board has informed us that, as described under Title 45 CFR Part 46 of the Code of Federal Regulations unidentifiable cadaver tissue does not constitute research in human subjects. Hence, the experiments performed in this report do not require their approval or waiver. Human corneal fibroblasts (HCFs) were isolated from cadaver corneas as previously described (Bernstein et al., 2007; Gillespie et al., 2017). HCFs were maintained with complete DMEM/F12 media (Gibco; Waltham, MA; Cat # 11330-032) with addition of 10% Fetal Bovine Serum (Atlanta Biologicals; Flowery Branch, GA) and 1X Antibiotic-Antimycotic (Sigma-Aldrich; St. Louis, MO). To prevent mycoplasma contamination, HCFs were also initially treated with Mycoplasma Removal Agent (Bio-Rad; Hercules, CA) after thawing from liquid nitrogen, and routinely with 10 μg/ml Plasmocin (Invivogen; San Diego, CA).

For all experiments, cells were passaged with TrypLE (Gibco; Waltham, MA), counted, and plated at either 2.5×10^4^ cells/well on glass coverslips in 24 well plates, 1×10^5^ cells per glassbottom dish (for live cell imaging; MatTek; Ashland, MA) or 1.2×10^6^ cells per 10 cm dish. Glass and plastic dishes were treated with 10 μg/ml Type I bovine collagen (Advanced BioMatrix; San Diego, CA) in PBS for >1hr at 37°C, then washed once before addition of HCFs. HCFs were then cultured in supplemented serum-free media (SSFM; 1 mM sodium pyruvate (Lonza; Basel, Switzerland), 2mM L-glutamine (Sigma-Aldrich; Waltham, MA), 1X MEM essential vitamin mixture (Gibco; Waltham, MA), 1X nonessential amino acid solution (Gibco; Waltham, MA), Insulin-Transferin-Selenium solution (Gibco; Waltham, MA), and 1X Antibiotic-Antimycotic solution, in DMEM/F12 media). TGFβ1 (R&D Systems; Minneapolis, MN) concentration for all relevant experiments was 2 ng/ml.

### Plasmid transfection and siRNA treatment

All transfections (2ug cDNA or 15 pmol siRNA) were carried out using the Lonza Nucleofector X-module using 1×10^6^-1.3×10^6^ cells per transfection. Transfections were conducted with Lonza’s P3 solution in 100 ul cuvettes using program EN-130. Cells were then plated on collagen-coated 10 cm dishes for biochemical analyses or MatTek glass bottom dishes.

### Immunocytochemistry

Cells were fixed using a protocol which is known to aid in the preservation of cytoskeletal morphology (Lunn et al., 1997; Smith-Clerc and Hinz, 2010). PHEM buffer (60mM PIPES, 25mM HEPES, 10mM EGTA, 2mM MgCl_2_, pH 6.9) and 3% PFA in PBS were warmed to 37°C before fixation. Cells were briefly rinsed in PHEM buffer twice and then fixed in 3% PFA for 15 min. Cells were then washed 3 times for 5 minutes in PBS at room temperature to remove PFA, and blocked in 10% goat serum (Thermo Fisher; Waltham, MA), 0.2% Triton X-100 in PBS for 1hr at room temperature. Cultures were then stained with a solution of primary antibody in 2% goat serum, 0.2% Triton X-100 in PBS for 2hr at room temperature. Primary antibodies and their concentrations were as follows: 1:1000 mouse-anti-α-SMA (Millipore Sigma; Burlington, MA; clone ASM-1/1A4), 1:300 rabbit-anti-Daam1 (Proteintech; Rosemont, IL; Cat # 14876-1-AP), and 1:300 mouse-anti-USP10 (Novus Biologicals; Centennial, CO; clone OTI2E1). For some experiments, Alexa Flour 555-conjugated phalloidin (Cytoskeleton Inc.; Denver, CO) was utilized as a counterstain at a concentration of 14 nM. After primary incubation, cells were again washed 3 times in PBS for 5 min per wash. Alexa Fluor 488-conjugated goat-anti-rabbit (1:500) and Alexa Fluor 647-conjuated goat-anti-mouse (1:800) secondary antibodies (Jackson Immunoresearch; West Grove, PA) were diluted in a solution consisting of 2% goat serum, 0.2% Triton X-100 in PBS and incubated on cultures for 1hr at room temp. Cultures were then washed 3 times in PBS, and coverslips were mounted with ProLong Gold mounting media with DAPI (Invitrogen) to glass slides. Following curing of mounting media, coverslips were sealed with clear nail polish and prepared for imagining.

### Proximity Ligation Assay

Duolink proximity ligation assay (Sigma) was carried out largely in congruence with manufacturer’s protocol, with some adjustments made after reaction optimization. Cultures were washed and fixed as described above. Before blocking, cells were permeabilized with 0.2% Triton X-100 in PBS for 5 minutes, and blocked with the kit blocking solution supplemented with an addition of 10% goat serum, 0.2% Triton X-100, and 2 ng/μL herring sperm DNA (ThermoFisher; Waltham, MA) for 1hr at 37°C. Cultures were incubated with primary antibodies diluted in the proprietary Antibody Diluent solution plus addition of 2% goat serum and 0.2% Triton X-100. For comparison of cultures either untreated or treated with 2ng/ml TGFβ1, 1:300 rabbit-anti-USP10 (Cell Signaling Technologies; Danvers, MA; Clone D7A5) and 1:300 mouse-anti-Daam1 (Novus Biologicals; Centennial, CO; Cat # NP_055807) were utilized. For comparison of Daam1-USP10 interaction to positive control (G3BP2), the following additional antibodies were used: 1:300 mouse-anti-USP10 (Novus Biologicals; Centennial, CO; clone OTI2E1) and 1:300 rabbit-anti-G3BP2 (Novus Biologicals; Cat # NBP1-82977). Cultures were incubated in primary antibody solution at 4°C overnight. The following day, cultures were washed 3 times with Duolink wash buffer A for 5 minutes per wash. Secondary antibody diluted as per manufacturer’s instructions (with addition of 2% goat serum and 0.2% Triton X-100) was applied and cultures incubated at 37°C for 1hr. Ligation and amplification/probe hybridization steps were carried out as per manufacturer’s instructions. Following amplification/probe hybridization step, cultures were washed with 1X Duolink wash buffer B for 10 minutes, and then twice with wash buffer A for 2 minutes each. HCFs were counterstained with 14nM Alexa Fluor-conjugated phalloidin in PBS for 1hr at room temperature, then washed twice with Wash Buffer A for 1 minute each. Cells were then briefly rinsed in 0.01X Wash Buffer B, and mounted as described above.

### Cell Imaging and Analysis

Epifluorescent imaging was carried out using a Nikon Eclipse N*i* upright fluorescence microscope with an Andor Zyla camera and NIS-Elements software (Nikon Instruments Inc.; Melville, NY). Live cell imaging was carried out using a Zeiss LSM780 confocal microscope, outfitted with a temperature and humidity-controlled chamber. Confocal images were acquired using ZEN software (Zeiss; Oberkochen, Germany). Image analysis, including immunoblots, was carried out using FIJI (ImageJ) software.

### qPCR

Cells were collected from culture dishes using trypsin. Total RNA was extracted from the corneal fibroblasts using the PureLink RNA kit (Invitrogen) according to manufacturer’s recommendations and was added as a template into a one-step multiplex qRT-PCR assay using Quanta qScript XLT ToughMix with ROX dye (VWR). For that, 1uL of total RNA was mixed with the reagent and primer-probe mixes for human Daam1, αv, β1, β5, and reference gene GAPDH in a 10uL reaction. The cycling parameters were as recommended by Quanta.

### Immunoblotting

HCFs grown on 10cm tissue-culture dishes were washed once in PBS and scraped from plates in RIPA buffer (0.1% SDS, 0.15 M NaCl, 0.5% sodium deoxycholate, 1% Triton X-100 in 0.05 M Tris (pH 6.8)) or a general solubilization buffer (2% SDS in 25 mM Tris (pH 8.8)) with 2mM PMSF and cOmplete protease inhibitor tablet. Lysates were then homogenized through a 26-gauge syringe 3 times and protein content measured by BCA (ThermoFisher; Waltham, MA). Samples were then diluted in 4X Sample Buffer (2% SDS, 10% glycerol, 1% β-mercaptoethanol, 12.5 mM EDTA, 0.02 % bromophenol blue in 50mM Tris (pH 6.8)), boiled at 95°C for 5 minutes, and loaded in pre-cast, 10% polyacrylamide gels (ThermoFisher; Waltham, MA). SDS-PAGE was carried out at 100V for 20 minutes for stacking phase and then 200V until completion.

Upon completion, polyacrylamide gels were transferred to Polyvinylidene fluoride (PVDF) membranes at 100V for 1hr. Membranes were then blocked for 1hr in tris-buffered saline with Tween-20 (TBST; Tris, sodium chloride, 0.1% Tween) with 5% goat serum and 1% bovine serum albumin (BSA) with gentle agitation. Following blocking, membranes were incubated in primary antibody solution consisting of 2% goat serum, 0.4% BSA in TBST, overnight at 4°C on a rocker. Antibodies used in this study are as follows: 1:1000 mouse-anti-USP10 (Novus Biologicals; Centennial, CO; clone OTI2E1), 1:1000 rabbit-anti-Daam1 (Proteintech; Rosemont, IL; Cat # 14876-1-AP), 1:2000 rabbit-anti-GAPDH (Cell Signaling Technologies, 14C10,), 1:2000 mouse-anti-α-tubulin (Cell Signaling Technologies, Danvers, MA; clone DM1A), 1:1000 mouse-anti-α-SMA (Millipore; Burlington, MA; Clone ASM-1), 1:2000 mouse-anti-β-actin (Cell Signaling Technologies; Danvers, MA; cat #: 3700), 1:1000 rabbit-anti-αv-integrin (Cell Signaling Technologies; Danvers, MA; cat #: 4711), 1:1000 rabbit-anti-α5-integrin (Cell Signaling Technologies; Danvers, MA; cat #: 4705), 1:1000 rabbit-anti-β1-integrin (Cell Signaling Technologies, Danvers, MA; cat #: 4706), 1:1000 rabbit-anti-β3-integrin (Cell Signaling Technologies, D7X3P-XP), and 1:1000 rabbit-anti-β5-integrin (Cell Signaling Technologies; Danvers, MA; clone D24A5). Subsequently, membranes were washed 3 times in TBST. Secondary antibodies were diluted in TBST with 2% goat serum, 0.4% BSA and incubated on membranes for 1hr at room temperature. Secondary antibodies were 1:3000 dilutions of horseradish peroxidase-conjugated goat-anti-mouse (Jackson Immunoresearch; West Grove, PA) and goat-anti-rabbit (EMD-Millipore; Burlington, MA). Membranes were washed 3 final times with TBST, and then incubated with chemiluminescent substrate (Thermo Fisher; Waltham, MA) briefly before imaging with a Chemidoc (Bio-Rad; Hercules, CA). Relative protein intensities were measured by ImageJ software.

### Stress Fiber Isolation

Stress Fiber Isolation protocol was based on previously published work by other groups (Katoh et al., 1998). Cells were washed briefly with chilled PBS and incubated in a solution of 2.5mM triethanolamine (pH 8.1) with cOmplete protease inhibitor and 2mM phenylmethylsulfonylfluoride (PMSF) for 20 minutes on ice and with gentle rocking. This solution was aspirated and replaced with 0.05% NP-40 in PBS with cOmplete protease inhibitor and 2mM PMSF for 10 min. Finally, this solution was aspirated and replaced with 0.5% Triton X-100 in PBS with cOmplete protease inhibitor and 2mM PMSF for 5 min. The last wash was aspirated and replaced with 200μL of fresh 0.5% Triton buffer, and the stress fibers scraped and collected in microcentrifuge tubes. The lysates were passed 3 times through 26-guage syringes and centrifuged at 100,000 x g for 1hr at 4°C. The supernatants were discarded and the pellets resuspended in solubilization buffer (2% SDS, 8M urea in 25mM Tris (pH 8.8) with 2mM PMSF and cOmplete protease inhibitor tablet). Final lysates were subjected to 1 freeze-thaw cycle before use in BCA and western blot, which aided the homogenization of cytoskeletal components. Lysates were then analyzed by immunoblot as described above.

### Ubiquant™ Ubiquitin Capture ELISA

Ubiquant™ analysis was carried out as described previously (Gillespie et al., 2017). HCFs were transfected with 150 pmol of siGLO nontargeting control siRNA (Dharmacon; Lafayette, CO), siDaam1, or siUSP10 (both: Santa Cruz Biotechnology; Dallas, TX) and incubated in SSFM on collagen-coated 10cm cell culture treated dishes for 24hrs. HCFs were then switched into SSFM with 5 μM MG132 and 10 μM chloroquine for 8hrs. Cultures were then scraped in RIPA buffer (as described above) plus complete protease inhibitor tablet (Roche; Basel, Swtizerland), PMSF (Thermo Fisher; Waltam, MA), 2mM NEM (Pierce; Waltham, MA) and 10 μg/ml PR-619 (Lifesensors; Malvern, PA). Protein content determined by BCA assay and samples were diluted in RIPA buffer to 0.4 μg/μL.

100 μL of lysate was added to each well (experiments carried out in triplicate) and incubated on a rocker at room temperature for 1hr. Wells were then washed 4 times with TBST and then incubated in 1x Blocking Buffer in PBS with a 1:10,000 dilutions of either β1- (Assay Biotechnology; San Francisco, CA; catalogue #: R12-2927) or β5-integrin (Assay Biotechnology; San Franscisco, CA; catalogue #: F-5) primary antibody for 1 hr at room temperature. Wells were washed again 4 times with TBST and incubated in the same buffer with a 1:15,000 dilution of goat anti-rabbit HRP-conjugated IgG for 1hr. Wells were washed a final 4 times, with 100 μL Developing Solution added to each well, and luminescence read by Epoch II plate reader and Gen5 software (BioTek Instruments; Winooski, VT).

### Biotinylated-FN Recycling Assay

HCFs were transfected (see above) with 150 pmol of either siGLO or siDaam1, then seeded in DMEM/F12 and 1% serum plus antibiotic-antimycotic solution. 24 hours post transfection, the cells were loaded with 10ug biotinylated-FN (Tang and Saito, 2017) for 3 hours. Cells were then passaged with trypsin and plated on 35mm glass bottom dishes in DMEM/F12 and 1% serum. After 48 hours, cells were washed 3 times for 30 minutes each prior to imaging with the following procedure: 1% PBSA in PHEM, 150mM Sodium Azide in PHEM, and 1:100 streptavidin-488 in PHEM. Images were analyzed using ImageJ’s 3D Object Counter plugin.

### Integrin Recycling Assay

HCFs were transfected (see above) with 150 pmol of either siGLO or siDaam1, then seeded in DMEM/F12 and 1% serum plus antibiotic-antimycotic solution 48 hrs post-transfection cells were blocked and treated with Ab against α5β1 (Novus, 2-52680) and αv (Cell Signaling 4711) at 10ug/ml for 30 minutes prior to cell surface stripping (0.2M acetic acid, 0.5M NaCl) for 30 sec. Cell were incubated for 90 min prior to incubation with 2° Ab-488 for 30 min. Live cells were imaged (Zeiss LSM 780 confocal) and analyzed using ImageJ’s Analyze Particles.

### Data and Statistical Analyses

All data was collected using Microsoft Excel software and graphs generated using GraphPad Prism software. Student’s unpaired, 2-tailed t-tests, logarithmic transformation, oneway ANOVA were calculated using Prism. Replicate number (n) refers to individual biological repeats (cell lines) derived from distinct human cadaver corneas. P-values: * *p*<0.05, ** *p*<0.01, and *** *p*<0.001, **** *p*<0.0001).

## Competing Interests

No competing interests declared.

## Acknowledgments

This work was supported by NIH-NEI R01 EY024942, NIH-NEI R01 EY030567, Merit Review Award (I01 BX005360) from the United States Department of Veteran’s Affairs, Biomedical Laboratory Research and Development Service, SUNY Upstate Start-up Funds, Unrestricted Grant to the Department of Ophthalmology & Visual Sciences from Research to Prevent Blindness, and The Lion’s District 20-Y.

## Notes

### Competing Interest Statement

The authors have declared no competing interest.

## References

Asano, Y., Ihn, H., Yamane, K., Jinnin, M. and Tamaki, K. (2006). Increased expression of integrin alphavbeta5 induces the myofibroblastic differentiation of dermal fibroblasts. Am J Pathol 168, 499–510.

Aspenstrom, P., Richnau, N. and Johansson, A. S. (2006). The diaphanous-related formin DAAM1 collaborates with the Rho GTPases RhoA and Cdc42, CIP4 and Src in regulating cell morphogenesis and actin dynamics. Exp Cell Res 312, 2180–94.

Basan, M., Elgeti, J., Hannezo, E., Rappel, W. J. and Levine, H. (2013). Alignment of cellular motility forces with tissue flow as a mechanism for efficient wound healing. Proc Natl Acad Sci U S A 110, 2452–9.

Benito-Jardon, M., Strohmeyer, N., Ortega-Sanchis, S., Bharadwaj, M., Moser, M., Muller, D. J., Fassler, R. and Costell, M. (2020). alphav-Class integrin binding to fibronectin is solely mediated by RGD and unaffected by an RGE mutation. J Cell Biol 219.

Benito-Jardon, M., Strohmeyer, N., Otega-Sanchis, S., Bharadwaj, M., Moser, M., Muller, D. J., Fassler, R. and Costell, M. (2021). Correction: alphav-Class integrin binding to fibronectin is solely mediated by RGD and unaffected by an RGE mutation. J Cell Biol 220.

Bernstein, A. M., Twining, S. S., Warejcka, D. J., Tall, E. and Masur, S. K. (2007). Urokinase receptor cleavage: a crucial step in fibroblast-to-myofibroblast differentiation. Mol Biol Cell 18, 2716–27.

Bharadwaj, M., Strohmeyer, N., Colo, G. P., Helenius, J., Beerenwinkel, N., Schiller, H. B., Fassler, R. and Muller, D. J. (2017). alphaV-class integrins exert dual roles on alpha5beta1 integrins to strengthen adhesion to fibronectin. Nat Commun 8, 14348.

Blanco-Mezquita, J. T., Hutcheon, A. E., Stepp, M. A. and Zieske, J. D. (2011). alphaVbeta6 integrin promotes corneal wound healing. Invest Ophthalmol Vis Sci 52, 8505–13.

Blanco-Mezquita, J. T., Hutcheon, A. E. and Zieske, J. D. (2013). Role of thrombospondin-1 in repair of penetrating corneal wounds. Invest Ophthalmol Vis Sci 54, 6262–8.

Bomberger, J. M., Barnaby, R. L. and Stanton, B. A. (2009). The deubiquitinating enzyme USP10 regulates the post-endocytic sorting of cystic fibrosis transmembrane conductance regulator in airway epithelial cells. J Biol Chem 284, 18778–89.

Boumil, E. F., Castro, N., Phillips, A. T., Chatterton, J. E., McCauley, S. M., Wolfson, A. D., Shmushkovich, T., Ridilla, M. and Bernstein, A. M. (2020). USP10 Targeted Self-Deliverable siRNA to Prevent Scarring in the Cornea. Mol Ther Nucleic Acids 21, 1029–1043.

Bridgewater, R. E., Norman, J. C. and Caswell, P. T. (2012). Integrin trafficking at a glance. J Cell Sci 125, 3695–701.

Bukowiecki, A., Hos, D., Cursiefen, C. and Eming, S. A. (2017). Wound-Healing Studies in Cornea and Skin: Parallels, Differences and Opportunities. Int J Mol Sci 18.

Castro, N., Gillespie, S. R. and Bernstein, A. M. (2019). Ex Vivo Corneal Organ Culture Model for Wound Healing Studies. J Vis Exp.

Chang, Y., Lau, W. L., Jo, H., Tsujino, K., Gewin, L., Reed, N. I., Atakilit, A., Nunes, A. C., DeGrado, W. F. and Sheppard, D. (2017). Pharmacologic Blockade of alphavbeta1 Integrin Ameliorates Renal Failure and Fibrosis In Vivo. J Am Soc Nephrol.

Corkins, M. E., Krneta-Stankic, V., Kloc, M., McCrea, P. D., Gladden, A. B. and Miller, R. K. (2019). Divergent roles of the Wnt/PCP Formin Daam1 in renal ciliogenesis. PLoS One 14, e0221698.

Danen, E. H. and Sonnenberg, A. (2003). Integrins in regulation of tissue development and function. J Pathol 200, 471–80.

Desmouliere, A., Redard, M., Darby, I. and Gabbiani, G. (1995). Apoptosis mediates the decrease in cellularity during the transition between granulation tissue and scar. Am J Pathol 146, 56–66.

Duperret, E. K., Natale, C. A., Monteleon, C., Dahal, A. and Ridky, T. W. (2016). The integrin alphav-TGFbeta signaling axis is necessary for epidermal proliferation during cutaneous wound healing. Cell Cycle, 1–10.

Esue, O., Harris, E. S., Higgs, H. N. and Wirtz, D. (2008). The filamentous actin cross-linking/bundling activity of mammalian formins. J Mol Biol 384, 324–34.

Gillespie, S. R., Tedesco, L. J., Wang, L. and Bernstein, A. M. (2017). The deubiquitylase USP10 regulates integrin beta1 and beta5 and fibrotic wound healing. J Cell Sci 130, 3481–3495.

Harris, E. S., Li, F. and Higgs, H. N. (2004). The mouse formin, FRLalpha, slows actin filament barbed end elongation, competes with capping protein, accelerates polymerization from monomers, and severs filaments. J Biol Chem 279, 20076–87.

Henderson, N. C., Arnold, T. D., Katamura, Y., Giacomini, M. M., Rodriguez, J. D., McCarty, J. H., Pellicoro, A., Raschperger, E., Betsholtz, C., Ruminski, P. G. et al. (2013). Targeting of alphav integrin identifies a core molecular pathway that regulates fibrosis in several organs. Nat Med 19, 1617–24.

Higgs, H. N. (2005). Formin proteins: a domain-based approach. Trends Biochem Sci 30, 342–53.

Higgs, H. N. and Peterson, K. J. (2005). Phylogenetic analysis of the formin homology 2 domain. Mol Biol Cell 16, 1–13.

Horan, G. S., Wood, S., Ona, V., Li, D. J., Lukashev, M. E., Weinreb, P. H., Simon, K. J., Hahm, K., Allaire, N. E., Rinaldi, N. J. et al. (2008). Partial inhibition of integrin alpha(v)beta6 prevents pulmonary fibrosis without exacerbating inflammation. Am J Respir Crit Care Med 177, 56–65.

Hsia, H. C., Nair, M. R. and Corbett, S. A. (2014). The fate of internalized alpha5 integrin is regulated by matrix-capable fibronectin. J Surg Res 191, 268–79.

Huttenlocher, A. and Horwitz, A. R. (2011). Integrins in cell migration. Cold Spring Harb Perspect Biol 3, a005074.

Huveneers, S., Truong, H., Fassler, R., Sonnenberg, A. and Danen, E. H. (2008). Binding of soluble fibronectin to integrin alpha5 beta1 - link to focal adhesion redistribution and contractile shape. J Cell Sci 121, 2452–62.

Jaiswal, R., Breitsprecher, D., Collins, A., Correa, I. R., Jr., Xu, M. Q. and Goode, B. L. (2013). The formin Daam1 and fascin directly collaborate to promote filopodia formation. Curr Biol 23, 1373–9.

Jonker, C. T. H., Galmes, R., Veenendaal, T., Ten Brink, C., van der Welle, R. E. N., Liv, N., de Rooij, J., Peden, A. A., van der Sluijs, P., Margadant, C. et al. (2018). Vps3 and Vps8 control integrin trafficking from early to recycling endosomes and regulate integrin-dependent functions. Nat Commun 9, 792.

Ju, R., Cirone, P., Lin, S., Griesbach, H., Slusarski, D. C. and Crews, C. M. (2010). Activation of the planar cell polarity formin DAAM1 leads to inhibition of endothelial cell proliferation, migration, and angiogenesis. Proc Natl Acad Sci U S A 107, 6906–11.

Katoh, K., Kano, Y., Masuda, M., Onishi, H. and Fujiwara, K. (1998). Isolation and contraction of the stress fiber. Mol Biol Cell 9, 1919–38.

Konigshoff, M., Balsara, N., Pfaff, E. M., Kramer, M., Chrobak, I., Seeger, W. and Eickelberg, O. (2008). Functional Wnt signaling is increased in idiopathic pulmonary fibrosis. PLoS One 3, e2142.

Lassance, L., Marino, G. K., Medeiros, C. S., Thangavadivel, S. and Wilson, S. E. (2018). Fibrocyte migration, differentiation and apoptosis during the corneal wound healing response to injury. Exp Eye Res 170, 177–187.

Liu, W., Sato, A., Khadka, D., Bharti, R., Diaz, H., Runnels, L. W. and Habas, R. (2008). Mechanism of activation of the Formin protein Daam1. Proc Natl Acad Sci U S A 105, 210–5.

Lobert, V. H. and Stenmark, H. (2010). Ubiquitination of alpha-integrin cytoplasmic tails. Commun Integr Biol 3, 583–5.

Lorenzo-Martin, E., Gallego-Munoz, P., Mar, S., Fernandez, I., Cidad, P. and Martinez-Garcia, M. C. (2019). Dynamic changes of the extracellular matrix during corneal wound healing. Exp Eye Res 186, 107704.

Lunn, K. F., Baas, P. W. and Duncan, I. D. (1997). Microtubule organization and stability in the oligodendrocyte. J Neurosci 17, 4921–32.

Lygoe, K. A., Norman, J. T., Marshall, J. F. and Lewis, M. P. (2004). AlphaV integrins play an important role in myofibroblast differentiation. Wound Repair Regen 12, 461–70.

Manichaikul, A., Wang, X. Q., Sun, L., Dupuis, J., Borczuk, A. C., Nguyen, J. N., Raghu, G., Hoffman, E. A., Onengut-Gumuscu, S., Farber, E. A. et al. (2017). Genome-wide association study of subclinical interstitial lung disease in MESA. Respir Res 18, 97.

Mao, Y. and Schwarzbauer, J. E. (2005). Fibronectin fibrillogenesis, a cell-mediated matrix assembly process. Matrix Biol 24, 389–99.

Massoudi, D., Malecaze, F. and Galiacy, S. D. (2016). Collagens and proteoglycans of the cornea: importance in transparency and visual disorders. Cell Tissue Res 363, 337–49.

Munger, J. S. and Sheppard, D. (2011). Cross talk among TGF-beta signaling pathways, integrins, and the extracellular matrix. Cold Spring Harb Perspect Biol 3, a005017.

Newman, D. R., Sills, W. S., Hanrahan, K., Ziegler, A., Tidd, K. M., Cook, E. and Sannes, P. L. (2016). Expression of WNT5A in Idiopathic Pulmonary Fibrosis and Its Control by TGF-beta and WNT7B in Human Lung Fibroblasts. J Histochem Cytochem 64, 99–111.

Nishimura, T., Ito, S., Saito, H., Hiver, S., Shigetomi, K., Ikenouchi, J. and Takeichi, M. (2016). DAAM1 stabilizes epithelial junctions by restraining WAVE complex-dependent lateral membrane motility. J Cell Biol 215, 559–573.

Pakshir, P. and Hinz, B. (2018). The big five in fibrosis: Macrophages, myofibroblasts, matrix, mechanics, and miscommunication. Matrix Biol 68-69, 81–93.

Pankov, R., Momchilova, A., Stefanova, N. and Yamada, K. M. (2019). Characterization of stitch adhesions: Fibronectin-containing cell-cell contacts formed by fibroblasts. Exp Cell Res 384, 111616.

Phillips, A. T., Boumil, E. F., Castro, N., Venkatesan, A., Gallo, E., Adams, J. J., Sidhu, S. S. and Bernstein, A. M. (2021). USP10 Promotes Fibronectin Recycling, Secretion, and Organization. Invest Ophthalmol Vis Sci 62, 15.

Reed, N. I., Jo, H., Chen, C., Tsujino, K., Arnold, T. D., DeGrado, W. F. and Sheppard, D. (2015). The alphavbeta1 integrin plays a critical in vivo role in tissue fibrosis. Sci Transl Med 7, 288ra79.

Rydell-Tormanen, K., Zhou, X. H., Hallgren, O., Einarsson, J., Eriksson, L., Andersson-Sjoland, A. and Westergren-Thorsson, G. (2016). Aberrant nonfibrotic parenchyma in idiopathic pulmonary fibrosis is correlated with decreased beta-catenin inhibition and increased Wnt5a/b interaction. Physiol Rep 4.

Saito, S., Tampe, B., Muller, G. A. and Zeisberg, M. (2015). Primary cilia modulate balance of canonical and non-canonical Wnt signaling responses in the injured kidney. Fibrogenesis Tissue Repair 8, 6.

Sandbo, N. and Dulin, N. (2011). Actin cytoskeleton in myofibroblast differentiation: ultrastructure defining form and driving function. Transl Res 158, 181–96.

Sarrazy, V., Koehler, A., Chow, M. L., Zimina, E., Li, C. X., Kato, H., Caldarone, C. A. and Hinz, B. (2014). Integrins alphavbeta5 and alphavbeta3 promote latent TGF-beta1 activation by human cardiac fibroblast contraction. Cardiovasc Res 102, 407–17.

Schiller, H. B. and Fassler, R. (2013). Mechanosensitivity and compositional dynamics of cell-matrix adhesions. EMBO Rep 14, 509–19.

Schmidt, S. and Friedl, P. (2010). Interstitial cell migration: integrin-dependent and alternative adhesion mechanisms. Cell Tissue Res 339, 83–92.

Schwarzbauer, J. E. and Sechler, J. L. (1999). Fibronectin fibrillogenesis: a paradigm for extracellular matrix assembly. Curr Opin Cell Biol 11, 622–7.

Shu, D. Y. and Lovicu, F. J. (2017). Myofibroblast transdifferentiation: The dark force in ocular wound healing and fibrosis. Prog Retin Eye Res 60, 44–65.

Smith-Clerc, J. and Hinz, B. (2010). Immunofluorescence detection of the cytoskeleton and extracellular matrix in tissue and cultured cells. Methods Mol Biol 611, 43–57.

Stepp, M. A., Zieske, J. D., Trinkaus-Randall, V., Kyne, B. M., Pal-Ghosh, S., Tadvalkar, G. and Pajoohesh-Ganji, A. (2014). Wounding the cornea to learn how it heals. Exp Eye Res 121C, 178–193.

Takayama, K. I., Suzuki, T., Fujimura, T., Takahashi, S. and Inoue, S. (2018). Association of USP10 with G3BP2 Inhibits p53 Signaling and Contributes to Poor Outcome in Prostate Cancer. Mol Cancer Res 16, 846–856.

Tang, J. and Saito, T. (2017). Human plasma fibronectin promotes proliferation and differentiation of odontoblast. J Appl Oral Sci 25, 299–309.

Thannickal, V. J., Lee, D. Y., White, E. S., Cui, Z., Larios, J. M., Chacon, R., Horowitz, J. C., Day, R. M. and Thomas, P. E. (2003). Myofibroblast differentiation by transforming growth factor-beta1 is dependent on cell adhesion and integrin signaling via focal adhesion kinase. J Biol Chem 278, 12384–9.

Wang, L., Pedroja, B. S., Meyers, E. E., Garcia, A. L., Twining, S. S. and Bernstein, A. M. (2012). Degradation of Internalized alphavbeta5 Integrin Is Controlled by uPAR Bound uPA: Effect on beta1 Integrin Activity and alpha-SMA Stress Fiber Assembly. PLoS One 7, e33915.

Whitcher, J. P., Srinivasan, M. and Upadhyay, M. P. (2001). Corneal blindness: a global perspective. Bull World Health Organ 79, 214–21.

Wierzbicka-Patynowski, I., Mao, Y. and Schwarzbauer, J. E. (2004). Analysis of fibronectin matrix assembly. Curr Protoc Cell Biol Chapter 10, Unit 10 12.

Wilson, S. E. (2012). Corneal myofibroblast biology and pathobiology: Generation, persistence, and transparency. Exp Eye Res 99, 78–88.

Wilson, S. E., Marino, G. K., Torricelli, A. A. M. and Medeiros, C. S. (2017). Injury and defective regeneration of the epithelial basement membrane in corneal fibrosis: A paradigm for fibrosis in other organs? Matrix Biol 64, 17–26.

Yanai, S., Wakayama, M., Nakayama, H., Shinozaki, M., Tsukuma, H., Tochigi, N., Nemoto, T., Saji, T. and Shibuya, K. (2017). Implication of overexpression of dishevelled-associated activator of morphogenesis 1 (Daam-1) for the pathogenesis of human Idiopathic Pulmonary Arterial Hypertension (IPAH). Diagn Pathol 12, 25.

Yang, C., Czech, L., Gerboth, S., Kojima, S., Scita, G. and Svitkina, T. (2007). Novel roles of formin mDia2 in lamellipodia and filopodia formation in motile cells. PLoS Biol 5, e317.

Yuan, J., Luo, K., Zhang, L., Cheville, J. C. and Lou, Z. (2010). USP10 regulates p53 localization and stability by deubiquitinating p53. Cell 140, 384–96.

Zatloukal, B., Kufferath, I., Thueringer, A., Landegren, U., Zatloukal, K. and Haybaeck, J. (2014). Sensitivity and specificity of in situ proximity ligation for protein interaction analysis in a model of steatohepatitis with Mallory-Denk bodies. PLoS One 9, e96690.

Zhang, H., Zhang, S. H., He, H. W., Zhang, C. X., Yu, D. K. and Shao, R. G. (2013). Downregulation of G3BPs inhibits the growth, migration and invasion of human lung carcinoma H1299 cells by suppressing the Src/FAK-associated signaling pathway. Cancer Gene Ther 20, 622–9.

Zhu, Y., Tian, Y., Du, J., Hu, Z., Yang, L., Liu, J. and Gu, L. (2012). Dvl2-dependent activation of Daam1 and RhoA regulates Wnt5a-induced breast cancer cell migration. PLoS One 7, e37823.

